# Groundwater redox dynamics across the terrestrial-aquatic interface of Lake Erie coastal ecosystems

**DOI:** 10.1101/2023.06.12.544684

**Authors:** Fausto Machado-Silva, Michael Weintraub, Nicholas Ward, Kennedy O. Doro, Peter J. Regier, Solomon Ehosioke, Shan Pushpajom Thomas, Roberta B. Peixoto, Leticia Sandoval, Inke Forbrich, Kenneth M. Kemner, Edward J. O’Loughlin, Lucie Setten, Trisha Spanbauer, Thomas B. Bridgeman, Teri O’Meara, Kenton A. Rod, Kaizad Patel, Nate G. McDowell, Ben P. Bond-Lamberty, J. Patrick Megonigal, Rich L. Rich, Vanessa L Bailey

## Abstract

Groundwater biogeochemistry in coastal areas is spatially and temporally dynamic because fluctuations in groundwater level may cause alternate redox between distinct hydrological conditions. Recent studies have proposed connections between biogeochemistry and large-scale hydrological processes, specifically focusing on the role of redox-active compounds in changing the oxidation state during flooding and draining events. While water saturation generally results in a shift of redox-active compounds from electron donors to acceptors, the specific mechanisms underlying the transition of groundwater between oxidizing and reducing conditions in response to water level fluctuations are uncertain. To determine the effects of groundwater levels on redox dynamics, we monitored groundwater redox potential across the terrestrial-aquatic interface in Lake Erie coastal areas throughout the high and low-water seasons. In contrast to previously observed responses to flooding in soils, our results revealed patterns of oxidizing redox potentials during high-water and reducing during low-water periods. Furthermore, short-term fluctuations in water table levels significantly impacted the redox potential of groundwater when dissolved oxygen increased, and redox dynamics displayed voltage hysteresis in most events. Based on these findings, we propose that for improved predictions of microbial functions and biogeochemical cycles, redox-informed models should incorporate the antagonistic changes in groundwater redox balance compared to soils and consider the time lags in redox fluctuations.

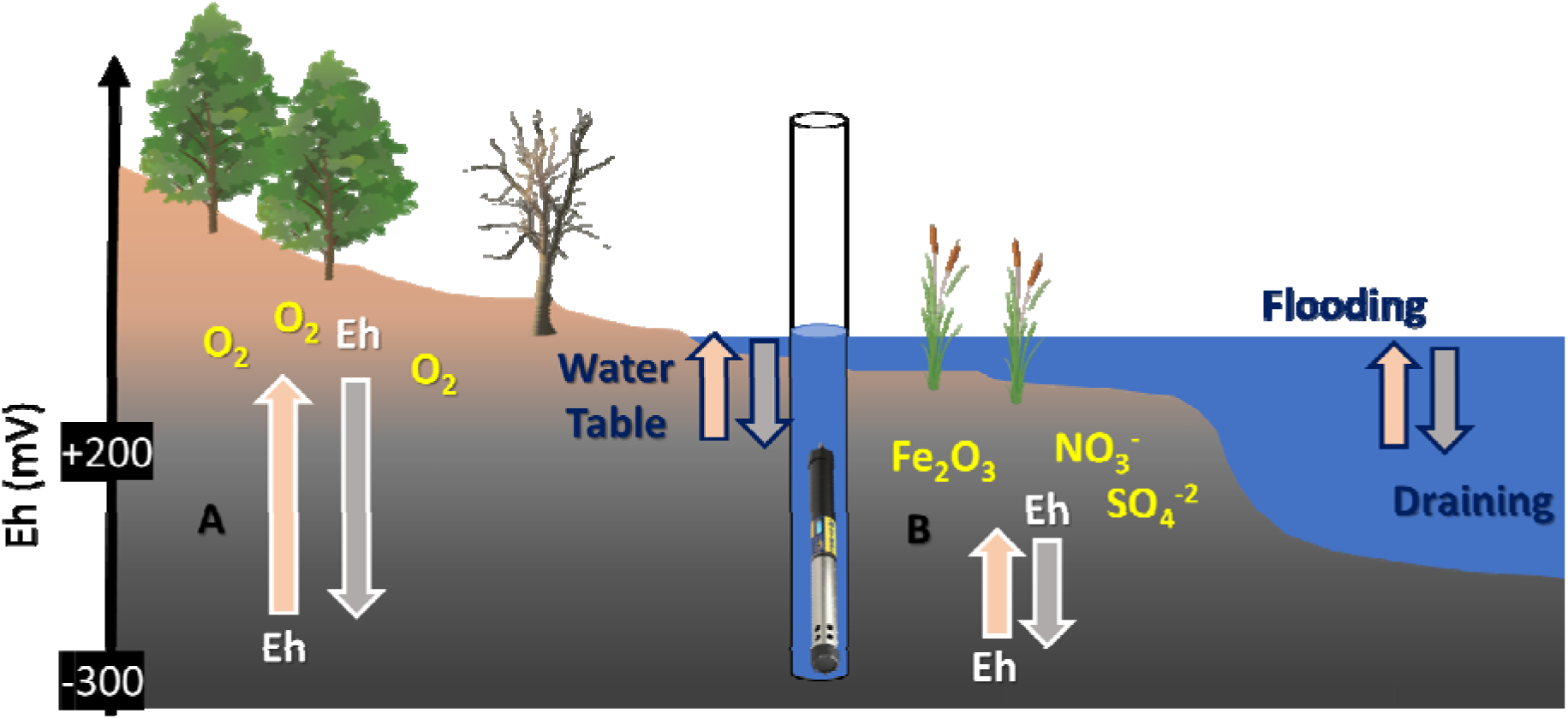
Graphical Abstract. Conceptual diagram of groundwater redox fluctuations in coastal ecosystems. Large redox fluctuations are derived by dissolved oxygen inputs and smaller more frequent redox fluctuations are led by redox sensitive species leaching from topsoil.

## 1. INTRODUCTION

Groundwater dynamics are intricately tied to the cycles of biogeochemical elements through the effects of water saturation on redox potential (Eh), the capability of chemical components in a solution to either oxidize or reduce other chemical components (Burgin and Loecke 2023). The interaction between surface water and groundwater commonly gives rise to mixing zones characterized by varying redox gradients, which can facilitate accelerated rates of carbon and nutrient cycling (Gómez-Gener et al. 2021). Groundwater Eh serves as a valuable proxy for assessing biogeochemical characteristics, predicting and understanding the predominance of organic matter cycling, and quantifying chemical fluxes from aquifers to coastal waters (Santos et al. 2021; Zhang and Planavsky 2020) and to the atmosphere (Fiedler and Sommer 2000; Olid et al. 2022). However, additional work is needed to determine the mechanisms and consequences of fluctuating groundwater levels on redox dynamics across the terrestrial-aquatic interface (Rosentreter et al. 2021; Yazbeck and Bohrer 2023; Zakem et al. 2020; Zhang and Planavsky 2020).

Coastal groundwater has been increasingly impacted by water table fluctuations in response to the intensification of the hydrological cycle, which may alter groundwater biogeochemistry (Burri et al. 2019). This motivates increasing demand for understanding and modeling the effects of water table fluctuations on groundwater biogeochemical processes (Pi et al. 2023; Wang and Chen 2021). Fluctuations of the groundwater can occur over multiple spatial and temporal scales due to patterns of infiltration and evapotranspiration that may vary according to precipitation patterns, temperature, hydraulic conductivity, freezing, and thawing (Condon et al. 2021; Gómez-Gener et al. 2021). Groundwater redox may vary due to changes in the concentration of redox-sensitive elements, which is dependent on aquifer water volume (Kim et al. 2020), the dissolution of solid elements present in the soil (Wang and Chen 2021) and entrapped gas of the pore space in the quasi-saturated zone (Klump et al. 2008; Kohfahl et al. 2009).

Groundwater and wetlands are often characterized by oxygen-depleted conditions and coastal aquifers’ redox potential exerts significant control over the transport of reduced solutes and nutrients into adjacent water bodies (Olid et al. 2022; Santos et al. 2021). Aquifer charging and discharging regimes can alter the redox state of groundwater and promote chemical reactions that affect concentrations of dissolved constituents. For instance, denitrification is a preferable microbial metabolic pathway in the absence of dissolved oxygen due to high energetic yields, removing nitrate and eventually producing nitrous oxide (Zhang et al. 2023). The biogeochemistry of iron and sulfur also plays a vital role in redoximorphic systems, mediating important transformations and reactions within these environments (Fan et al. 2017; Frohne et al. 2011; Fulda et al. 2013). Even though it yields the lowest energy, methanogenesis can prevail as the primary pathway for organic matter degradation in deep soil and groundwater, especially when other electron donors have been exhausted, resulting in the production of methane that is subsequently released in wetland areas or neighboring water bodies (Fiedler and Sommer 2000; Olid et al. 2022; Peixoto et al. 2015; Rosentreter et al. 2021; Santos et al. 2021).

Fluctuations in redox dynamics have been observed in a variety of soils under flooding regimes. For instance, flooded soils are often reported with lower redox than dry soils in floodplains, (Aeppli et al. 2022; Jia et al. 2020; Megonigal et al. 1993), coastal marshes (Noyce et al. 2023), and agricultural systems (Honma et al. 2016; Wang et al. 2020), highlighting the impact of dynamic hydrological changes. Experiments also confirm the influence of flooding and draining on the transition between reducing and oxidizing phases (Pi et al. 2023; Rezanezhad et al. 2014; Zhao et al. 2023). These findings support the recently proposed biogeobattery concept, based on alternating charge and discharge of electrons, providing energy for microbial processes in environments under varying hydrological regimes (Kappler et al. 2021; Peiffer et al. 2021; Zhao et al. 2023). In particular, it is expected that oxygen becomes limited during flooding, leading to a reducing phase characterized by a negative charge in the system. Conversely, during draining, increased gas diffusion promotes an oxidizing phase, causing the release of previously stored electrons and exposing the system to fluctuating redox conditions. Fluctuations in groundwater and soil chemistry under changing hydrological regimes may determine ecosystem redox state and the dominance of microbial processes defined by the dominant redox couples.

However, despite the growing recognition of the influence of hydrological conditions on groundwater redox fluctuations, there is still limited understanding of how groundwater redox dynamics vary in response to specific hydrological regimes (Meng et al. 2021; Wanner et al. 2019) (Meng et al. 2021). Furthermore, variations in dissolved oxygen and redox during water recharge periods remain poorly understood (Kumar and Riyazuddin 2012; Yu et al. 2022). Previous studies highlight the complex nature of groundwater redox dynamics, emphasizing the need for further research and continuous monitoring to unravel the underlying mechanisms and better understand the role of hydrological conditions in shaping subsurface biogeochemistry(Zhang and Furman 2021). Addressing these knowledge gaps is crucial for a comprehensive understanding of subsurface biogeochemistry and its connections to large-scale hydrological dynamics

Therefore, the objective of this study was to investigate the dynamics of groundwater redox potential in coastal ecosystems during the natural transition from high to low waters. We hypothesized: (1) there would be higher variability in groundwater redox potential at transitional areas from uplands to wetlands due to high water table variability; (2) groundwater redox would potential increase during draining as the limitation to oxygen diffusion in highly water-saturated soil is alleviated. We tested these hypotheses by evaluating groundwater levels, water chemistry parameters, and redox potential along three elevation gradients across the Lake Erie terrestrial-aquatic interface.

## 2. METHODS

### 2.1. Study area

Our study was conducted in three sites located along the Ohio shoreline of Lake Erie, spanning from approximately 41°39’4.57“N and 83°14’39.16”W to 41°21’53.14“N and 82°30’16.78”W (Figure 1a). We addressed groundwater in a transversal gradient of the terrestrial-aquatic interface, including wetlands, transitional areas, and uplands (Figure 1b), situated in three tributaries: Crane Creek (CRC), Portage River (PTR), and Old Woman Creek (OWC). The CRC site is situated in a drowned-river mouth wetland complex that spans more than 11 km from east to west and 4 km from north to south, encompassing the Ottawa National Wildlife Refuge (ONWR) and other protected areas. The PTR site is located near the confluence of the Portage River on the west side of the Little Portage River, adjacent to the Little Portage River Wildlife Area. The OWC site is situated within an Ohio state nature reserve, known as Old Woman Creek, which is a component of NOAA’s National Estuarine Research Reserve (NERR) network. The OWC estuary, which bathes the coastal areas, is located behind barrier beaches, and its opening is influenced by climate and environmental variations.

**Figure 1.**
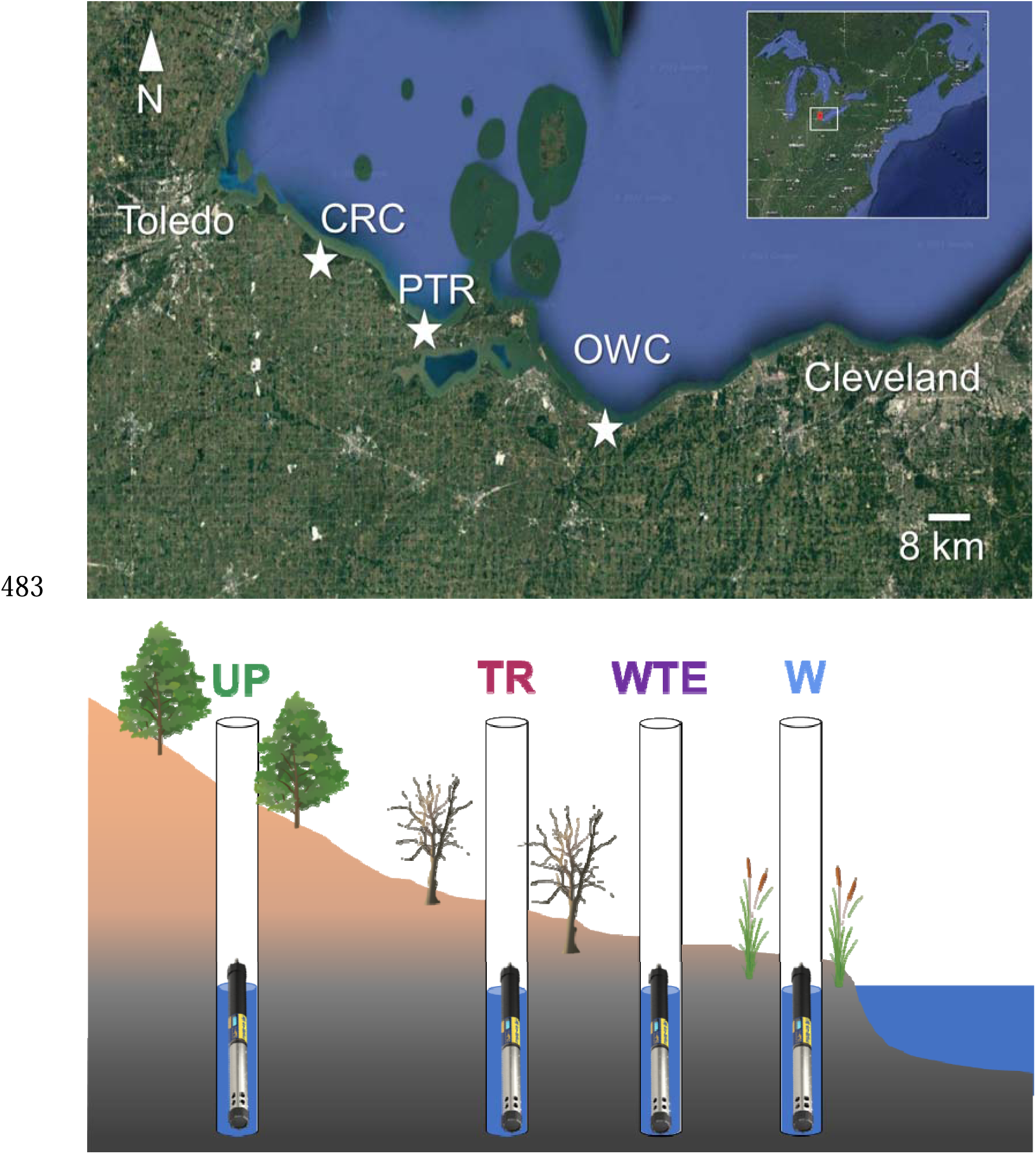
A. Lake Erie coastal ecosystems and B. Conceptual diagram of the terrestrial aquatic interface with groundwater multiparameter sondes in uplands, transition, wetland transition edge and wetlands.

These coastal forests and marshes are unique and ecologically important habitats located at the confluence of rivers or streams flowing into Lake Erie (Herdendorf 1992) and are characterized by surficial deposits primarily composed of clayey till formation from the Quaternary period (USGS 2023). Fluctuations in water levels within the wetland are primarily driven by interannual and short-term variations in Lake Erie’s water levels (NOAA 2023), while the influence of watershed inputs can either amplify or mitigate the effects of these changes, particularly after storm events (Farhadzadeh et al. 2017; Kowalski et al. 2009; NOAA 2023; Swatridge et al. 2022).

### 2.2. Groundwater and soil measurements

Each study area was equipped with three groundwater wells, one in each of three zones: upland, upland-wetland transition, and wetland. Wells were made of 4.2 cm diameter PVC tubes with 2.5 mm slots at 1 cm intervals on the bottom 1.52 m of the well (ESP Supply). The remaining length of the well was solid sealed at the surface with bentonite. In case of 1 m wells 1 m deep with 2.5 mm slots at 1 cm interval across the entire belowground length (ESP Supply). Piezometer depths varied from 4.9 to 5.1 m at uplands, 1.3 to 2.5 m at transition, and from 0.3 to 1.3 m at wetlands and was 0.6 m at the wetland-transition edge. Each well contained one vented multiparameter water quality sonde (In situ, Aquatroll 600) measuring water oxidation-redox potential (ORP), pH, temperature, conductivity, dissolved oxygen, and depth. Sensors were equipped with wipers to minimize fouling of sensor heads and calibrated per manufacturer protocols during maintenance visits and measured at 15-min intervals. In addition, soil ORP was measured monthly in the field from March 16^th^ to November 30^th^, 2022 using electrode based redox probes (SWAP Instruments, Netherlands) at 5, 10 and 20 cm from soil surface.

Both groundwater and soil ORP were corrected to the standard hydrogen electrode before analysis by applying the following equation derived from empirical measurements of commonly used reference electrodes versus the standard hydrogen electrode at different temperatures (ISO 11271:2022, APHA Method 2580).

Eh (mV) = ORP (mV) –0.718*T +224.41

Where Eh represents a measure of soil redox potential measured in mV referred to a Standard Hydrogen Electrode, which comprises the H^+^/H_2_ redox couple, having a standard potential of 0 mV. ORP is the oxidative-reduction potential measured in the field with an inert platinum electrode and the Ag/AgCl 3M KCl electrode, and is the soil temperature measured in °C.

### 2.3. Electrical resistivity tomography (ERT)

ERT data were collected across the three sites with SuperSting R8 and 84 electrodes switch box (Advanced Geosciences Inc., Austin, TX), using the dipole-dipole electrode configuration and 1 m spacing between the electrodes (Loke 1999). The data was collected in automatic mode which automatically record resistivity data using a preprogrammed command file and the distributed Swift automatic multi-electrode system (AGIUSA 2005). The acquired resistivity data was inverted with the AGI Earth Imager 2D using smoothness constrained inversion method and then converted to conductivity. Finally, the Earth Imager was used to trim the conductivity profiles to the desired depth for correlation with other soil properties.

### 2.4. Statistical analysis

We determined median and interquartile range, of groundwater level, groundwater quality parameters, and soil Eh, for the entire study period. To evaluate the normality of the groundwater level and Eh, we employed the Anderson-Darling test (Stephens, 1974), which is best suited for large sample sizes and places focus on the tails of the distribution. To capture significant differences in groundwater levels and Eh between zones and sites, we used the Friedman test with repeated measures in date and time in pairwise comparisons using data from simultaneous measurements across all sites and zones and used the site-zone combination as a fixed factor. We also computed the effect size to estimate the magnitude of the observed differences. Additionally, to look for significant variations between site-zones, we performed post hoc multiple comparisons using the Wilcoxon rank sum test. We used the Bonferroni correction for the overall Type I error rate.

We determined median and interquartile range of soil Eh. To evaluate the normality of the soil Eh at different depths, which was manually taken providing discrete measurements at smaller sample size than groundwater monitoring, we employed the Shapiro-Wilk test that provides a more accurate assessment to small sample sizes being more powerful compared to other normality tests (Razali & Wah, 2011). To capture significant differences in soil Eh between zones, sites, and depths, we applied a linear mixed-effects model with a random intercept for the date of measurement to estimate fixed effects of depth and random effects of date. We then conducted pairwise comparisons using Tukey-Kramer criterion to assess significant differences between specific depth categories within sites and zones. These statistical methods provided a robust analysis of the differences in soil profile Eh across depths by comparing median values. The use of mixed model analysis allowed us to account for within-subject correlation and incorporate random effects, while pairwise comparisons allowed us to assess differences between specific depth categories.

To test weather groundwater Eh is influenced by groundwater dissolved oxygen, we extracted from time series events of oxygen above 0.01 mg/L (our lower limit of detection) and applied linear regression models to assess the prediction of a linear relationship. We also estimated Pearson coefficients of determination to quantify the proportion of the total variation in the dependent variable groundwater Eh that can be explained by the independent variable groundwater dissolved oxygen.

## 3. RESULTS

### 3.1. Spatial differences across the terrestrial aquatic interface

#### 3.1.1. Groundwater level and redox potential variations

Figure 2 shows a box plot summarizing the spatial variation of groundwater level and Eh across the study area, highlighting the different conditions between the three sites (CRC, OWC, and PTR) and three zones (UP, TR, W). Groundwater level is shown as negative when below the soil surface and positive when above. As expected, we observed a gradient of deeper groundwater levels in the uplands and shallower in the wetlands, except for the wetland-transition edge (WTE) at OWC-WTE, whose groundwater level was below that of the transition (TR), which was counter to the expected pattern. Overall, the uplands had the lowest mean groundwater levels and the largest range. In particular, PTR-UP had the lowest mean and median, while CRCUP fell to the lowest level, and had the largest range and interquartile range. By contrast, the highest mean groundwater levels were in the wetlands, in which OWC-W shows a mean at soil surface and a maximum above the soil surface (Figure 1a). Groundwater Eh mean, and median values were higher in TR than W in the three sites. Although mean levels were higher in UP than TR in both CRC and PTR, there was a large overlap between zones, and in the case of CRC-UP, the median was even lower than CRC-TR. The values range from a low of –245 mV for PTR-W to a high of 662 mV for OWC-WTE. The CRC site generally has negative Eh values, with the lowest mean (and median) at the upland. PTR-UP has the highest mean (and median), while OWC-WTE has the largest interquartile range (Figure 1b).

**Figure 2.**
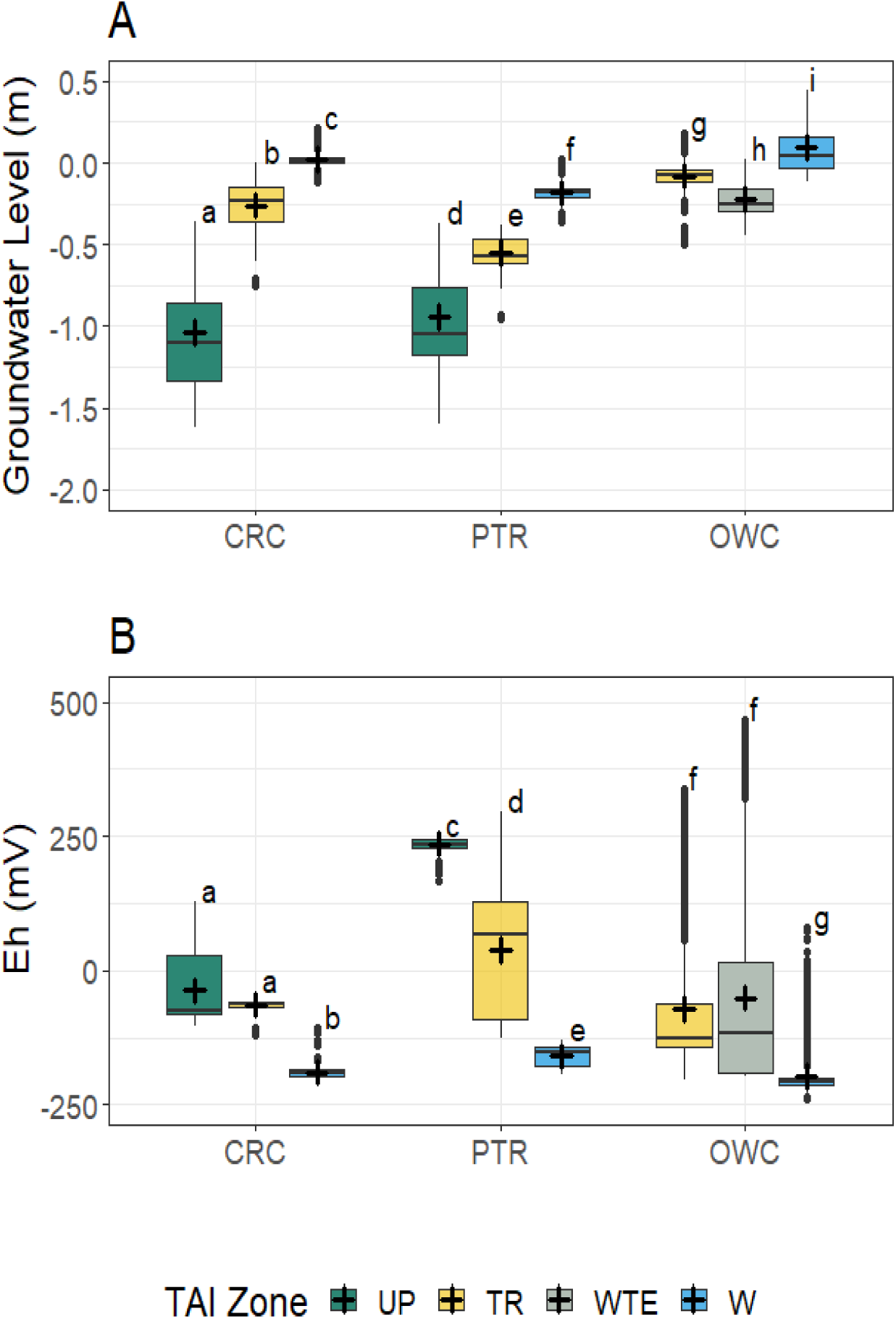
Groundwater level (m) and redox potential (Eh, mV) in the terrestrial-aquatic interface (TAI) zones of coastal ecosystems of Lake Erie. Coastal ecosystem sites are Crane Creek (CRC), Portage River (PTR), and Old Woman Creek (OWC) and zones of the TAI are Upland (UP), Transition (TR), Wetland-Transition Edge (WTE), and Wetland (W). Crosses show mean, bars median, boxes the interquartile range, and whiskers the 95% range. of Different letters indicate significant differences between site-zones (P<0.001).

#### 3.1.2. Groundwater vs soil redox potential (Eh)

Our analysis highlights significant disparities between soil Eh (redox potential) and groundwater Eh, with no overlap in the interquartile ranges (Table 1). The most pronounced differences were in the upland zones, particularly at the CRC site. In contrast, wetland areas show less apparent differences than uplands and transition but exhibit distinct interquartile ranges. Generally, soil Eh values are positive, while groundwater Eh values are predominantly negative. However, transitional areas such as PTR and OWC display negative values only in the first interquartile range (Table 1).

**Table 1.** Comparison between soil and groundwater redox potential (Eh, mV) in the terrestrialaquatic interface (TAI) zones of coastal ecosystems of Lake Erie. Zones of the TAI are Upland (UP), Transition (TR), and Wetland (W) and sites are Crane Creek (CRC), Portage River (PTR), and Old Woman Creek (OWC). Data is expressed as median (interquartile range).

| Zone | Site | Soil Eh (mV) | Groundwater Eh (mV) |
| --- | --- | --- | --- |
| UP | CRC | 569 (596 to 550) | -99 (50 to -120) |
| UP | PTR | 590 (613 to 563) | 224 (230 to 190) |
| UP | OWC | 572 (597 to 534) |  |
| TR | CRC | 390 (493 to 328) | -60 (-44 to -89) |
| TR | PTR | 577 (601 to 495) | 152 (256 to -19) |
| TR | OWC | 531 (558 to 498) | 109 (429 to -131) |
| WTE | OWC |  | -102 (192 to -138) |
| W | CRC | 410 (474 to 340) | -178 (-111 to -191) |
| W | PTR | 472 (507 to 428) | -170 (-153 to -190) |
| W | OWC | 111 (143 to 88) | -202 (-167 to -223) |

The zones of the terrestrial aquatic interface displayed distinct Eh values in the soil profile (Fig. 3; Linear mixed effects model, p < 0.05). For instance, we observed that: (1) uplands displayed the highest soil Eh mean and median and relatively less variable values with a slight increase from 5 to 20 cm depth; (2) mean soil Eh in transitional areas was between the values for uplands and wetlands in OWC and PTR, but not in CRC, which had a higher range. In addition, CRC-TR and PTR-TR showed a similar pattern of soil Eh decrease from 5 cm to 10 cm and an increase from 10 cm to 20 cm, and OWC-TR showed a very stable soil Eh with depth; (3) wetlands displayed lower soil Eh values and greater variation in the profile across the sites, which ranged between a minimum at a depth of 20 cm in OWC-W and a maximum at a depth of 10 cm in PTR-W. Interestingly, wetland soils had contrasting Eh profiles between sites. For example, while at CRC-W Eh increased with increasing depth, Eh decreased with depth at OWC-W. In CRC-W and OWC-W, the change from 5 cm to 10 cm is more prominent than the change from 10 cm to 20 cm. PTR wetland shows a slight increase in Eh from 5 cm to 10 cm and a decrease from 10 cm to 20 cm. In particular, the pairwise comparison indicated significant differences across depths 5, 10, and 20 cm performed within site and zones, excepting for transition at OWC and PTR (p<0.05, Figure 3).

**Figure 3.**
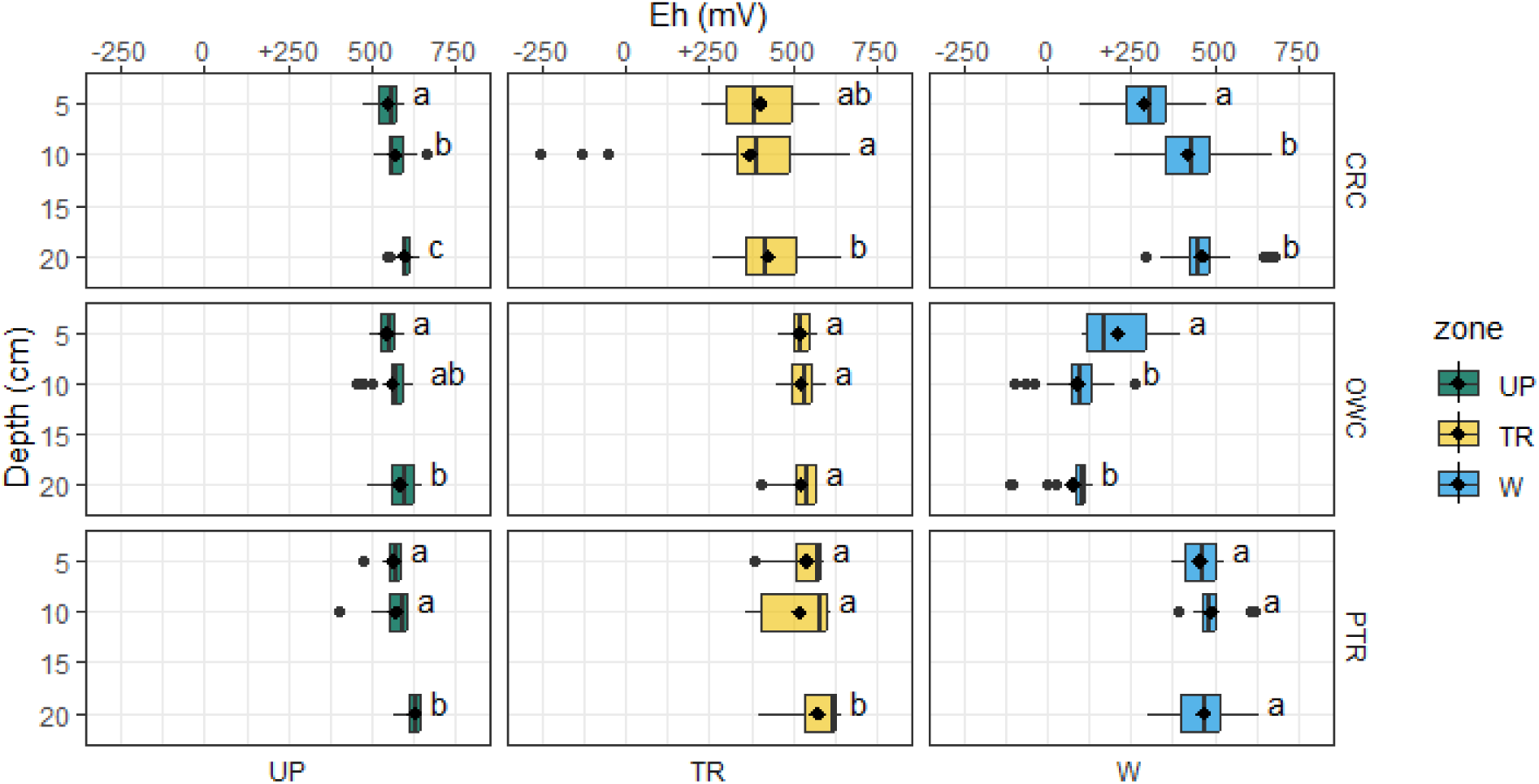
Soil redox potential (Eh Soil, mV) profile (depth, cm) in the terrestrial-aquatic interface (TAI) zones of coastal ecosystems of Lake Erie. Coastal ecosystem sites are Crane Creek (CRC), Portage River (PTR), and Old Woman Creek (OWC) and zones of the TAI are Upland (UP), Transition (TR), and Wetland (W). Bars show median, interquartile range, and 95% range. Dots show mean and values above 95% range. Different letters indicate statistical differences between depths within sites and zones (p<0.05).

### 3.2. Temporal dynamics across the terrestrial aquatic interface

#### 3.2.1. Seasonal trends in groundwater level and redox potential

A seasonal trend was observed at the Lake Erie terrestrial-aquatic interface (TAI) for both groundwater level and groundwater redox potential (Fig. 4). Groundwater levels exhibited variable and heterogeneous dynamics across the TAI. Generally, the uplands displayed a consistent downward trend in groundwater levels from spring through autumn, but CRC-UP had a more prominent decline than PTR-UP. The transition areas exhibited diverse variations over time. In CRC-TR, groundwater levels oscillated throughout the study period and trended down starting in June, falling below detection in September. In PTR-TR, there was a continuous decline in groundwater level, similar to the decrease observed in the uplands. In OWC, fluctuations in the groundwater level occurred, with the groundwater level reaching positive values at different times until July when it dropped below the detection limit. The wetland areas all tended to decline over the study. In CRC-W, the groundwater level was below detection in July. We observed three instances of more rapid increases in groundwater levels from below the detection to positive values. The groundwater level in PTR-W exhibited minor oscillations and declined from August onwards, reaching a level lower than the detection limit in September. At OWC-W, the groundwater remained above the ground surface level for most of the study period. OWC-WTE exhibited similar groundwater variability but was approximately 0.4 meters below OWC-W, on average (Fig. 4A).

**Figure 4.**
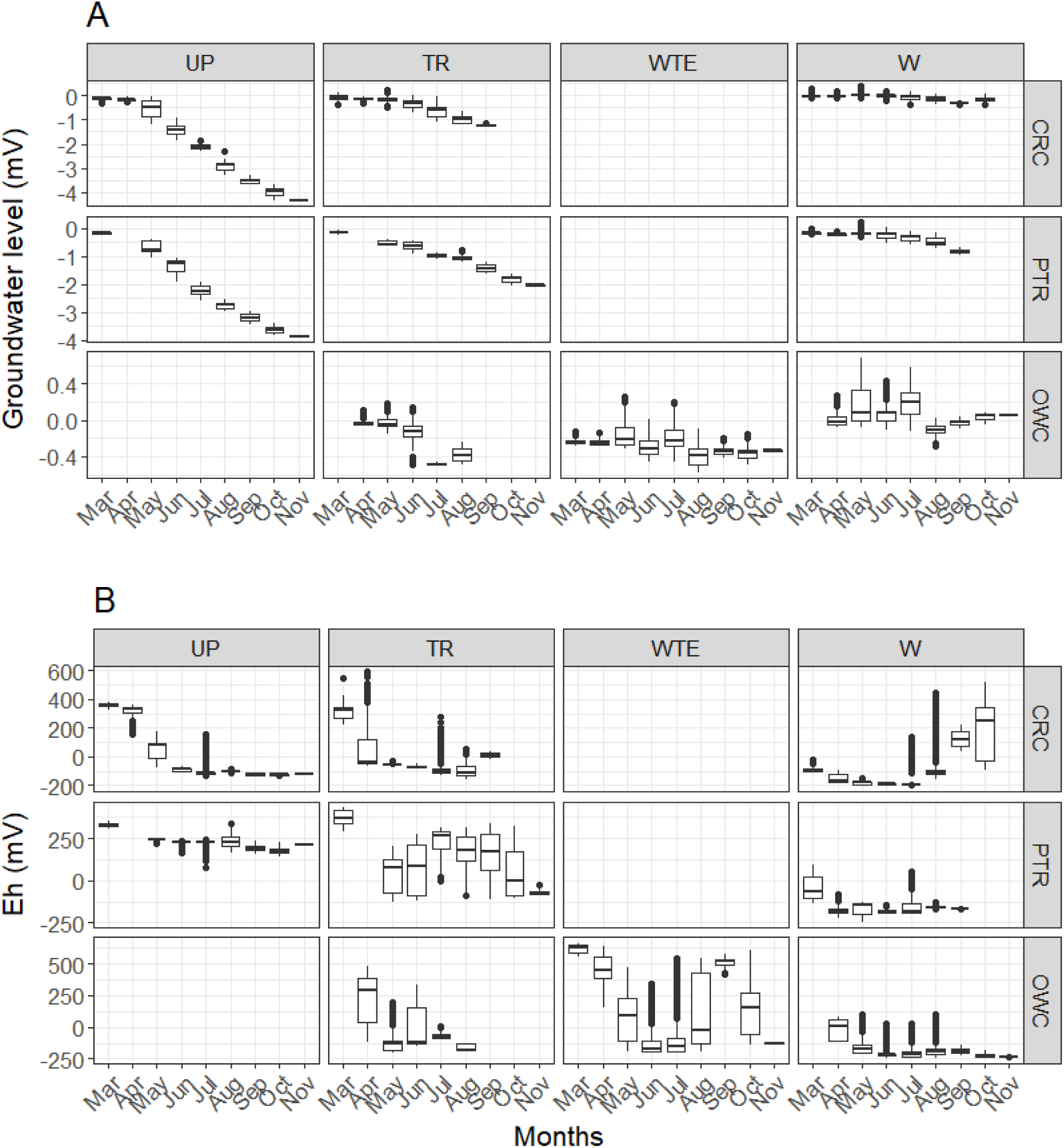
Seasonal patterns of groundwater level (m) in Lake Erie coastal areas. Graphs show zones of the terrestrial-aquatic interface Upland (UP), Transition (TR), Wetland Transition Edge (WTE), and Wetland (W) in sitesCrane Creek (CRC), Portage River (PTR), and Old Woman Creek (OWC).

Water table fluctuations showed high heterogeneity during the study period (Figure 5). For example, the water table of uplands exhibited a significant negative trend from high water table values close to the ground surface (0 m) at the beginning of monitoring to low values at the end of the survey and some intramonthly fluctuations. Water table at transitional areas in CRC and OWC displayed high intramonthly variability, but PTR showed a marked trend similar to the uplands. On wetlands, the water table displayed high intramonthly variability and water table variability at OWC WTE was very similar to OWC W.

**Figure 5.**
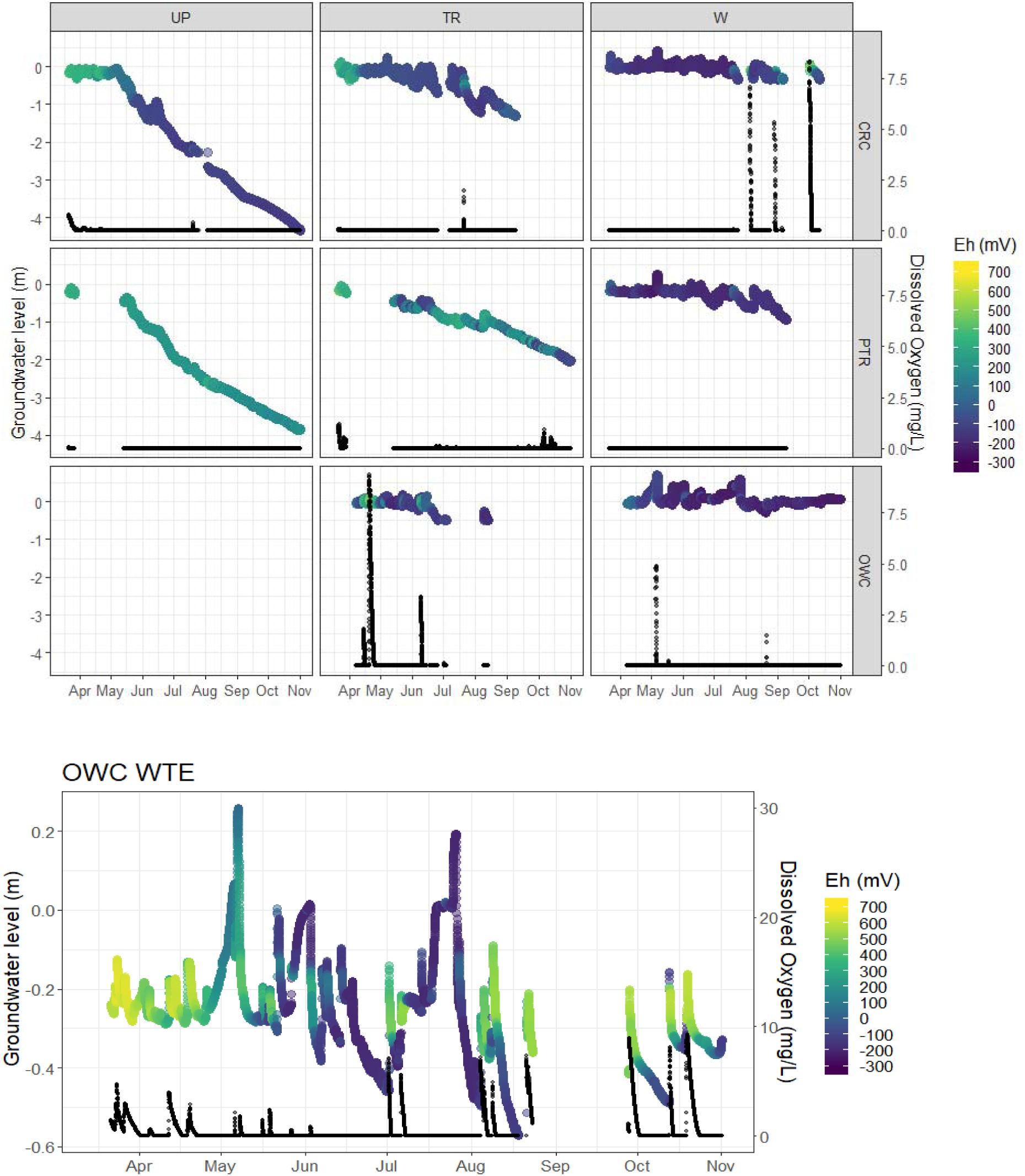
Dynamics of groundwater level (m), dissolved oxygen (mg/L), and redox potential (Eh, mV) in Lake Erie coastal areas. Graphs show zones of the terrestrial-aquatic interface Upland (UP), Transition (TR), Wetland Transition Edge (WTE), and Wetland (W) in sitesCrane Creek (CRC), Portage River (PTR), and Old Woman Creek (OWC).

In addition to water table fluctuations, groundwater redox potential dynamics reveal complex seasonality across the TAI gradient (Figure 5). Overall, the uplands and transition areas had the highest Eh in March and April. Groundwater Eh in CRC-UP declined prominently from March to November, although there was substantial variability in March-May. Groundwater Eh in PTRUP had a decreasing trend from March to May, then remaining stable until the end of the series with minor variability in September. In CRC-TR, groundwater Eh was highest in March, then declined until August, while September showed a median value approximately 100 mV higher than August (Figure 5). In PTR-TR, groundwater exhibited pronounced oscillations throughout the study period. Furthermore, a decrease in Eh from March to May, an increase in the median from May to June, and a reduction from June to November were observed. OWC WTE displayed the highest variation in Eh throughout the study period, with significant oscillations and values exceeding 600 mV. The median decreased from March to June, increased from June to September, and decreased from September to November. The wetlands generally exhibited less variable Eh values, mostly below 0 mV, except for CRC W, which showed higher deals starting from August, coinciding with events of elevated Eh values (Figure 4B).

#### 3.2.2. Short term dynamics of groundwater level, dissolved oxygen, and redox potential

We observed substantial variability in groundwater level fluctuations, redox potential, and dissolved oxygen concentrations at the terrestrial-aquatic interface. The most notable changes in redox potential occurred concomitantly with rising groundwater levels and subsequent oxygenation from surface water inputs (Figure 5). In the upland and transition areas in the spring, particularly in March and April, elevated redox potential (above 340 mV) coincided with oxygenated floods. Moreover, the transition areas experienced considerable groundwater level fluctuations that did not correspond with substantial redox potential or dissolved oxygen changes (Figure 5A).

Wetland areas displayed frequent water table increases from below –0.3 m to above surface level that were unrelated to oxygenation or significant shifts in redox potential (Figure 5A). Within the CRC-W site, we identified three episodes of rapid water table rise surpassing the soil surface. These events coincided with heightened levels of oxygenation and a more pronounced variation in redox potential, primarily occurring after August when the water table level fell below the detection limit. Specifically, in the OWC-WTE, we documented 19 flood events characterized by a marked increase from undetectable to up to 10 mg/L in dissolved oxygen levels and a subsequent rise in redox potential from below to above 0 mV. Nevertheless, flood events displayed lower redox variability and minimal or no oxidation (Figure 5B).

To investigate the influence of dissolved oxygen (DO) on groundwater redox potential, we extracted redox values from periods of oxygenation and tested linear relationships between them. There was a consistent positive association between dissolved oxygen and redox potential in spite of some variable Eh responses. Linear correlations between dissolved oxygen concentrations (from times when it was above detection) and Eh were significant, with coefficients of determination ranging from 0.91 to <1 (p<0.001, Figure 6).

**Figure 6.**
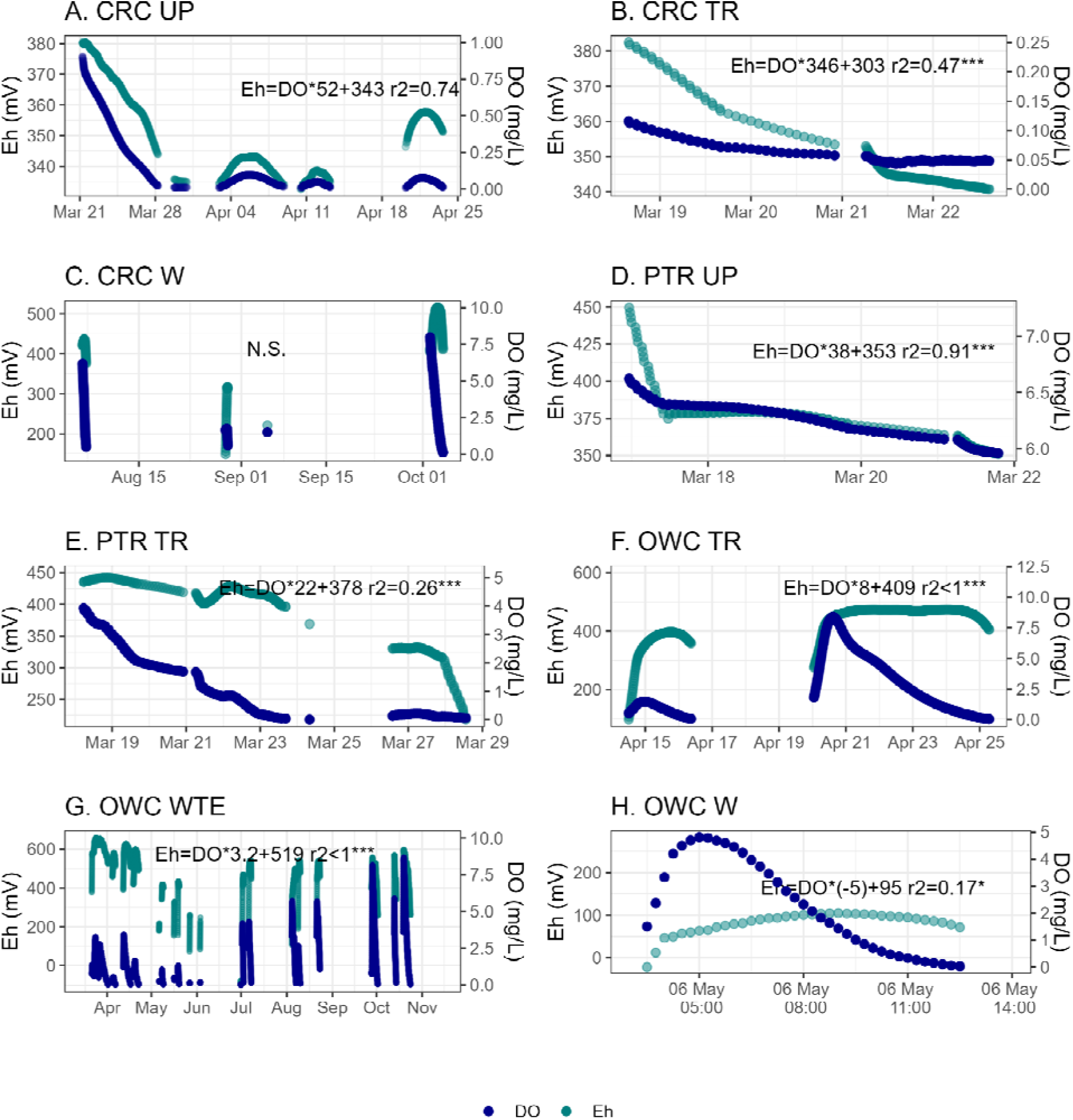
Redox potential (Eh, mV) *versus* dissolved oxygen (DO, mg/L) in groundwater of Lake Erie coastal zones. Graphs show Crane Creek (CRC), Portage River (PTR), and Old Woman Creek (OWC), and zones within sites Upland (UP), Transition (TR), Wetland Transition Edge (WTE), and Wetland (W).

In addition, to investigate sources of variability of the relationship between Eh and DO we graphed Eh *versus* DO and shaded from light to dark in chronological order to visualize the pattern of the parameter over the course of a water table rise. Then, we notice a remarkable delay in the changes of Eh concerning variations in DO concentrations during both the oxygenation and the oxygen consumption phases, revealing the presence of a counterclockwise hysteresis loop (Figure 7).

**Figure 7.**
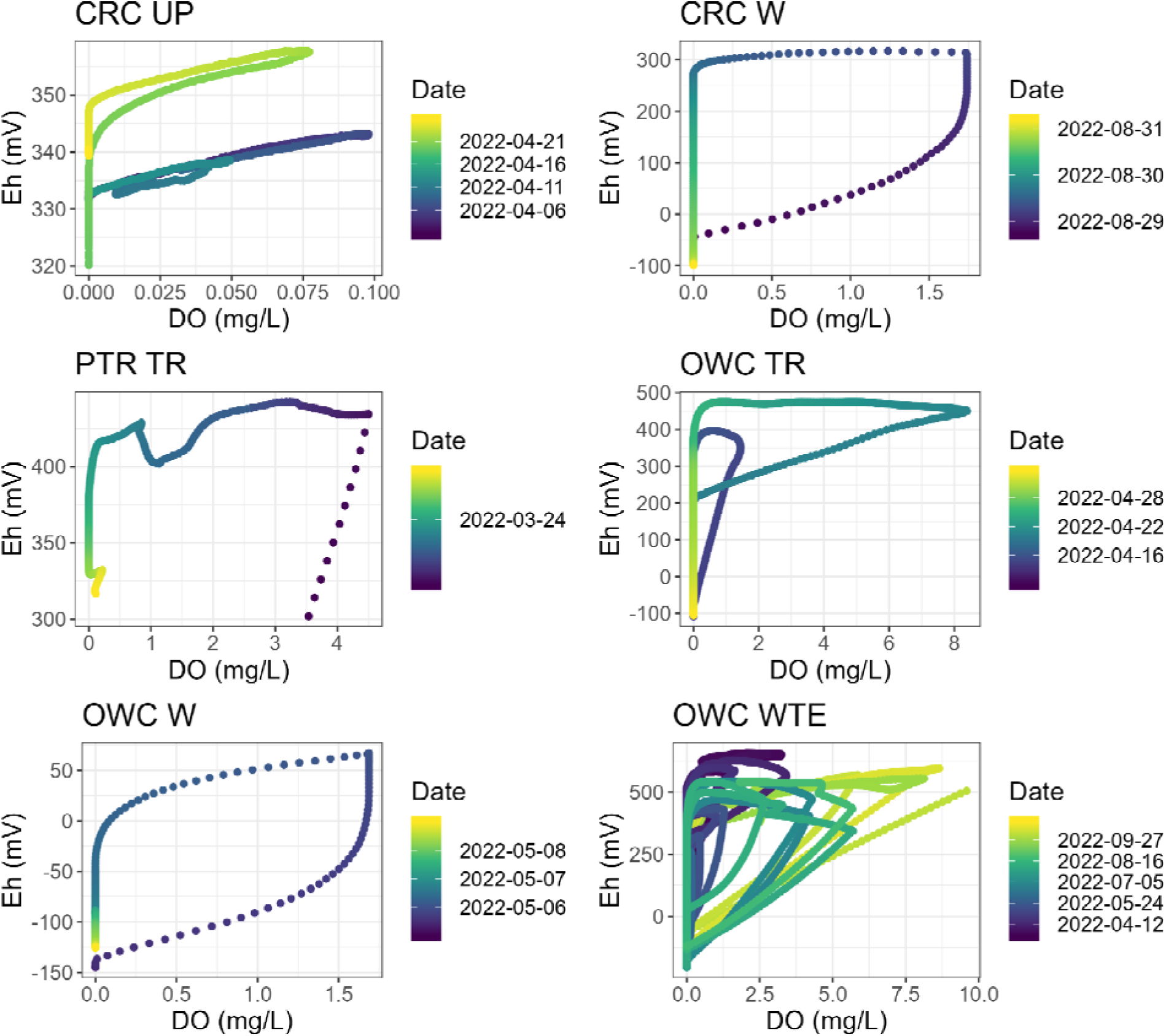
Redox potential (Eh, mV) hysteresis regarding dissolved oxygen (DO, mg/L) variability. Eh varies with DO counterclockwise and dates are displayed from dark to light colors.

### 3.3. Groundwater level and water quality across sites and zones

Heterogeneous patterns across sites and zones highlight variations in groundwater properties in terrestrial aquatic interface zones. Groundwater levels varied the most in the uplands, then in the transitional areas, and the least during the wetlands. The OWC-WTE varied in Eh the most, followed by the transition zones, while the wetlands showed the lowest variation (Table 2). Eh maximums had little impact on average dissolved oxygen concentrations, which remained near to zero. Groundwater pH was around neutral, but somewhat more basic in CRC-UP and slightly more acidic in PTR-UP. PTR-TR has the highest specific conductivity, followed by PTR-TR and CRC-UP. OWC, specifically the OWC-WTE, had the lowest mean conductivity. Groundwater temperature varied significantly in the wetlands and OWC-WTE, and was more consistent and stable in the uplands, with some intermediate variance in the transitions.

**Table 2.**
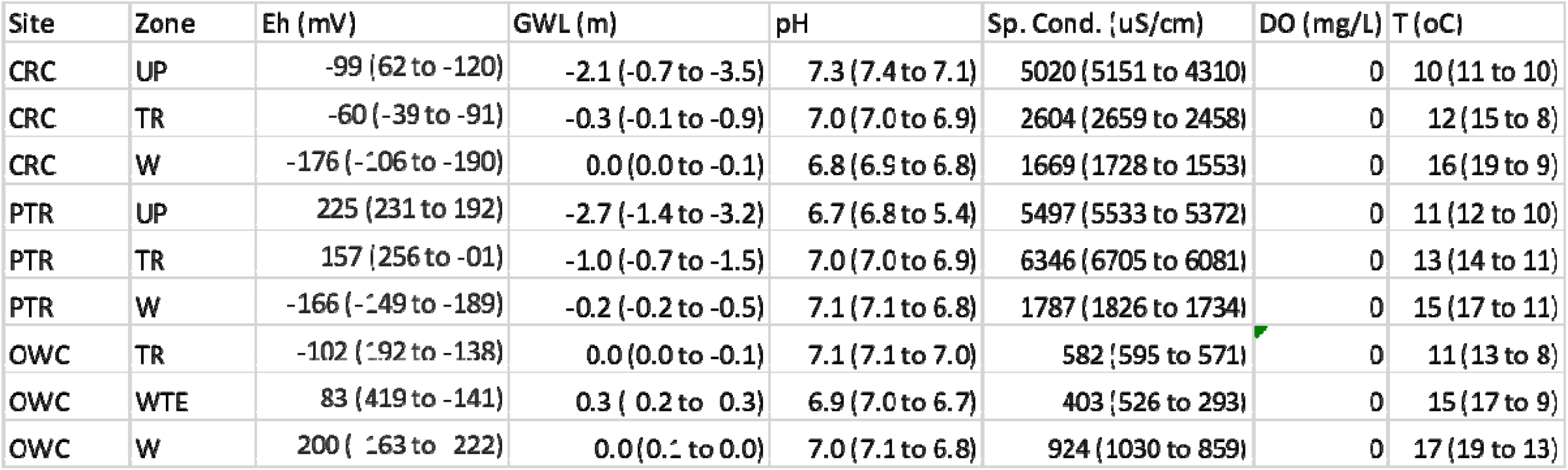
Summary of groundwater level and water quality parameters across the sites (Crane Creek – CRC, Portage River – PTR, and Old Woman Creek – OWC), and zones of the terrestrial aquatic interface (Upland – UP, transition –TR, and wetlands –W). Groundwater levels are reference of the soil surface in meters (m), and groundwater quality parameters are: redox potential (Eh, mV), dissolved oxygen (DO, mg L^-1^), pH, specific conductivity (Sp. Cond., µS cm^-1^), and temperature (°C).

### 3.4. Electrical Resistivity Tomography (ERT)

In this study, we utilized ERT to assess potential vertical and lateral connections in groundwater. The conductivity profiles obtained from ERT data provided insights into the water movement within the soil. Specifically, at the PTR upland site, the conductivity profile indicated water reaching the soil surface (topsoil), as evidenced by high conductivity values above mS/m, which indicated high water content (Figure 8A). Furthermore, the conductivity profile at the OWCWTE site revealed the influence of the stream on the hyporheic zone (bottom), with high conductivity values exceeding 80 mS/m observed from the surface to a depth of 3.25 m below the surface. These conductivity profiles offer valuable information regarding the extent and pathways of water movement in the subsurface, highlighting the potential for vertical and lateral connections in groundwater.

**Figure 8.**
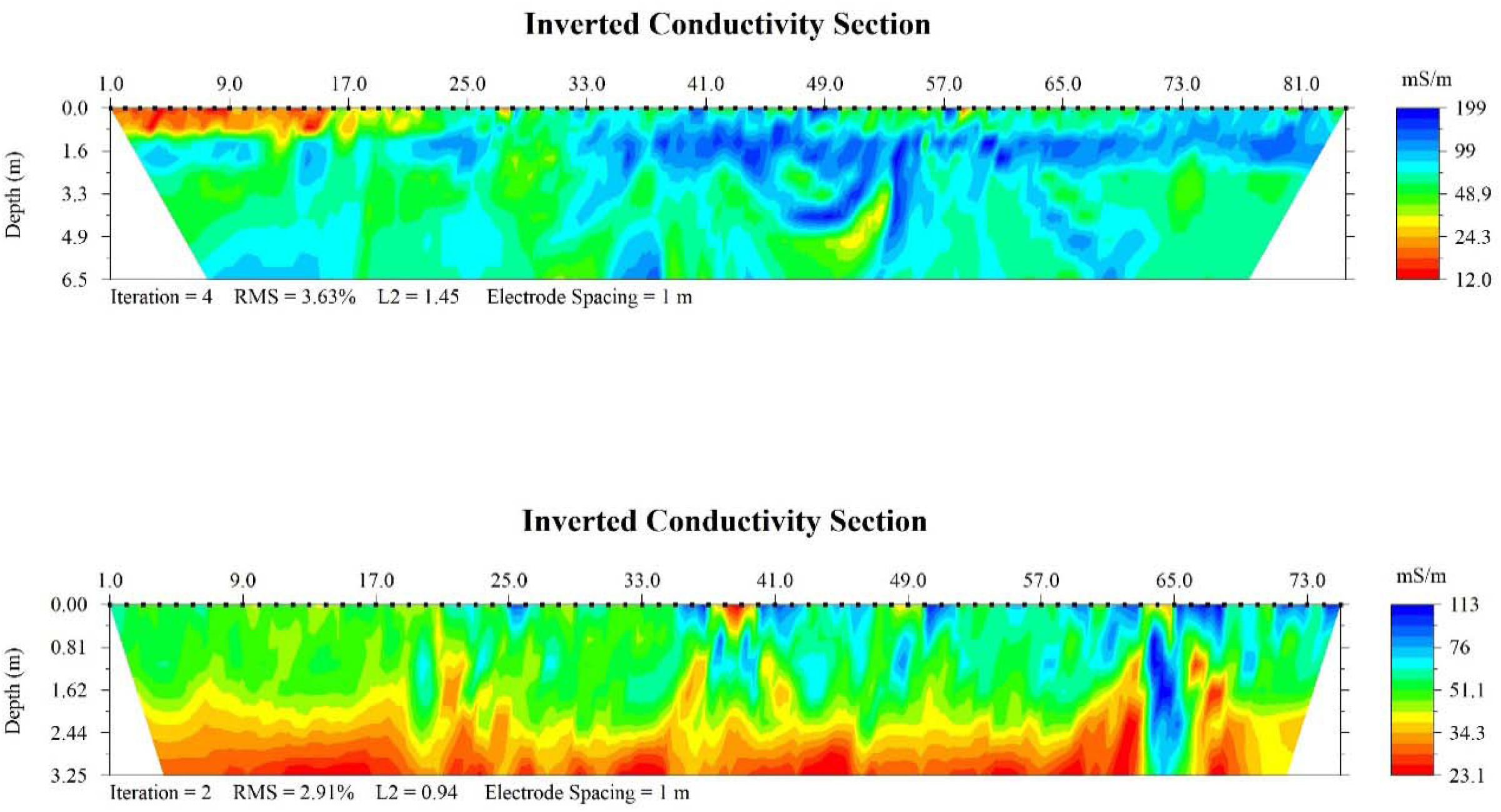
Conductivity profiles (mS/cm) indicates water achieving soil surface (top graph, e.g., PTR-UP profile) and the influence of stream into the hyporheic zone (bottom graph, e.g., OWC-WTE).

## 4. DISCUSSION

### 4.1. Groundwater levels and redox potential

We hypothesized that a decline in groundwater levels would lead to an increase in groundwater redox potential, assuming enhanced oxygen diffusion into drained soil. However, patterns of groundwater levels and redox potential did not follow this prediction. Instead, our results indicate that during the high-water season, the redox potential of groundwater is strongly influenced by the surface soil, suggesting that oxidized redox-sensitive element consumption occurs at slower rates compared to their turnover. Conversely, lower redox potential during drainage, despite a greater unsaturated soil volume, suggests that the deep soil plays a substantial role in determining groundwater redox potential, with electron acceptor depletion occurring rapidly. These results emphasize the complex interplay between surface soil and deep soil in influencing groundwater redox potential.

The significant impact of topsoil on the redox potential of groundwater stems from (1) the contrasting redox conditions in groundwater between high and low-water seasons (Figure 5) and (2) the conductivity profiles indicating groundwater links with the soil surface (Figure 8). Soil is generally characterized by abundant electron acceptors, which creates a more oxidizing environment (Zhang and Furman 2021) while groundwater tends to be reduced and anoxic (Fiedler and Sommer 2000). In periods of high water table level, groundwater may become hydrologically connected to surface soil, which enables the groundwater to receive oxygen or other oxidizing elements, leading to an increase in its reduction potential and elevating the redox potential of the groundwater (Kumar and Riyazuddin 2012). In addition, entrapped oxygen in soil pores may induce groundwater oxygenation during increased water table levels (Klump et al. 2008; Kohfahl et al. 2009). By contrast, during a decrease in the water table, the groundwater can become isolated, leading to a decline in redox potential likely caused by microbial consumption of electron acceptors (Fulda et al. 2013; O’Connor et al. 2018). Persistent reduced condition in groundwater were previously associated with the dominance of lateral connectivity in isolated aquifers (Richards et al. 2022). Therefore, our observations indicate the presence of an electron charging-discharging mechanism driven by the presence or absence of a hydrological connection between groundwater and surface soils, which in turn is controlled by flood conditions.

### 4.2. Redox potential depth and lateral variability

The concept of the redox ladder suggests a predictable reduction of electron acceptors with increasing depth (Burgin and Loecke 2023; Zhang and Furman 2021). Thus, the soil surface is commonly characterized as more oxidizing than the deeper soil, which exhibits more reducing conditions. Redox potential also varies with topography in coastal areas, with lower values typically observed in wetland environments (Noyce et al. 2023; Yu et al. 2006) and higher values in upland areas (Neogi et al. 2020; Yu et al. 2006). Our results were consistent with this pattern: we observed higher redox potential (Eh) in the soil compared to groundwater, and wetlands exhibited persistent reducing conditions and were rarely oxygenated (Table 1, Fig. 2). Of particular interest was the Old Woman Creek (OWC) wetland-transition edge (WTE), which displayed higher Eh values compared to the adjacent transition and wetland zones (Figure 2). We speculate that this due to the proximity to a stream 1 m away (Figure 8). The relatively high redox of the stream may have influenced the redox conditions of the surrounding areas (Gómez-Gener et al. 2021; Nimick et al. 2011), facilitating the interconnection between the hyporheic zone and groundwater and leading to elevated Eh values during flooding (Figure 5). In contrast, we found relatively consistent soil Eh values across the different zones at OWC (Table 2). However, it should be noted that in the case of OWC-W at depths of 10 and 20 cm, there were variations where the groundwater frequently rose to this level (Figure 3).

### 4.3. Redox potential seasonality

Despite numerous studies of variations in redox potential and dissolved oxygen with fluctuations in groundwater levels, we still lack a comprehensive understanding of the complex interactions and underlying mechanisms driving this variability. For instance, in a fractured bedrock aquifer in southwestern Ontario, Canada, researchers observed slight increases in redox potential and dissolved oxygen concentration during the low-water season (Wanner et al. 2019), as we had initially predicted for this study. Between high– and low-water intervals in a tidal channel in the USA, higher redox potential was distinguished due to oxygen inflows compared to both high– water and low-water periods (Yu et al. 2022). Intra-monthly variability in redox potential was found to be larger than seasonal variability in a riverbank environment in northeast China (Meng et al. 2021), highlighting the dynamic nature of redox processes. Similarly, studies in India have shown higher redox potential in shallow groundwater during high-water post-monsoon seasons (Kumar and Riyazuddin 2012). Similarly, studies in India have shown higher redox potential in shallow groundwater during high-water post-monsoon seasons.

Our study confirmed significant seasonal variations in groundwater levels in upland areas, which showed the highest range of redox potential variability. Despite some fluctuations, higher average values of Eh and dissolved oxygen (above 0 mV and 0 mg/L respectively) were observed in the spring than later in the growing season, indicating relatively more oxidizing conditions. In contrast, wetlands had a narrower range of water table variability and exhibited a consistently lower and more stable redox potential, typically below –100 mV. Groundwater redox potential was more heterogeneous in transitional areas, with a wide spectrum of values, except for CRC, which showed greater stability and was more similar to the uplands. Notably, OWC-WTE exhibited substantial variation in redox potential despite low groundwater level variability, possibly due to its connection to a nearby stream through the hyporheic zone (Gómez-Gener et al. 2021). These findings contribute to our understanding of seasonal variations in groundwater redox potential across different landscape zones and emphasize the influence of hydrological connectivity for groundwater redox dynamics.

### 4.4. Mechanism underlying short term variability

Generally, sudden increases in groundwater dissolved oxygen concentrations indicate short periods of vertical connectivity, even in environments dominated by lateral connectivity (Gómez-Gener et al. 2021; Regier et al. 2021). We observed fluctuations in groundwater levels from late winter to spring that were associated with increased oxygenation in the uplands and transitional areas (Figures 4 and 5). In particular, 19 oxygenation events occurred at the interface between the transition zone and the wetland at the OWC site (OWC-WTE). Interestingly, these oxygenation events were relatively evenly distributed throughout the study period and did not show clear seasonality.

Furthermore, during late autumn, we observed low water levels, with the water table falling below the detection limit in several study sites. However, there were also discrete events characterized by high groundwater and increased dissolved oxygen concentrations (Figure 5) (Farhadzadeh et al. 2017; Kowalski et al. 2009; NOAA 2023; Swatridge et al. 2022). For instance, the Toledo Water Level Station in Lake Erie recorded a 1-meter decline and subsequent rebound in lake depth from September 27 to October 3 (Figure S1). Notably, at the CRC-W site, we observed three events of water table oscillating from below detection limit to above soil surface simultaneously with groundwater Eh and DO peaks (Figure 5).

Then, Finally, groundwater redox potential (Eh) and dissolved oxygen levels were positively correlated in some fluctuations (Figure 6). However, we also observed complex behavior characterized by voltage hysteresis in charge transfer processes, with different redox potentials during charge and discharge (Figure 7). This hysteresis phenomenon can be observed when the response of water chemistry exhibits a delay in performing a loop-like behavior due to the difference between the rising and declining limb in fluctuating water levels (Indivero et al. 2021). Our findings suggest that the redox behavior is not solely determined by external conditions, such as oxygen availability, but also shows inertness over certain ranges of conditions before responding more strongly when reaching critical levels, such as oxygenation (Figure 7). This distinct response curve highlights the existence of two alternative stable states, with different redox dynamics, on the upper and lower branches (Scheffer et al. 2001).

## 5. Conclusion

Overall, these observations emphasize the complexity of groundwater redox dynamics and the importance of considering fluctuating water levels in understanding the interplay between redox potential, dissolved oxygen, and external conditions. Integrating redox processes into models poses a challenge but holds the potential to enhance the accuracy of simulations related to greenhouse gas emissions in dynamic coastal areas with hydrological and biogeochemical variations. Microbially driven biogeochemical reactions can use heterotrophic or autotrophic pathways, employ different redox-sensitive elements at varying rates, and are sources of atmospheric carbon dioxide, methane, and nitrous oxide, contributing to global warming (Ward et al. 2020; Zakem et al. 2020). However, groundwater can exhibit multiple simultaneous redox conditions and concurrent biogeochemical transformations under coexisting redox conditions (Li et al. 2023; Zuo et al. 2021). (Burgin and Loecke 2023; Megonigal and Rabenhorst 2013; Peiffer et al. 2021). (McMahon and Chapelle 2008). To develop accurate redox-informed models, it is essential to establish a theoretically grounded framework that can parameterize the diverse microbial metabolisms operating simultaneously (Zakem et al. 2020).

## AUTHOR INFORMATION

### Corresponding Author

*Fausto Machado-Silva, PhD.

Department of Environmental Sciences, University of Toledo.

3050 W. Towerview Blvd., Bowman-Oddy Laboratories, Room 3001A, Toledo, OH, 43606. Phones: Office: 419.530.2353 and Mobile: 419.344.9569

### Author Contributions

The manuscript was written through contributions of all authors.

## ACKNOWLEDGMENT

This research was found by the U.S. Department of Energy and acknowledge the University of Toledo group and collaborators

## Supporting Information

**Figure S1.**
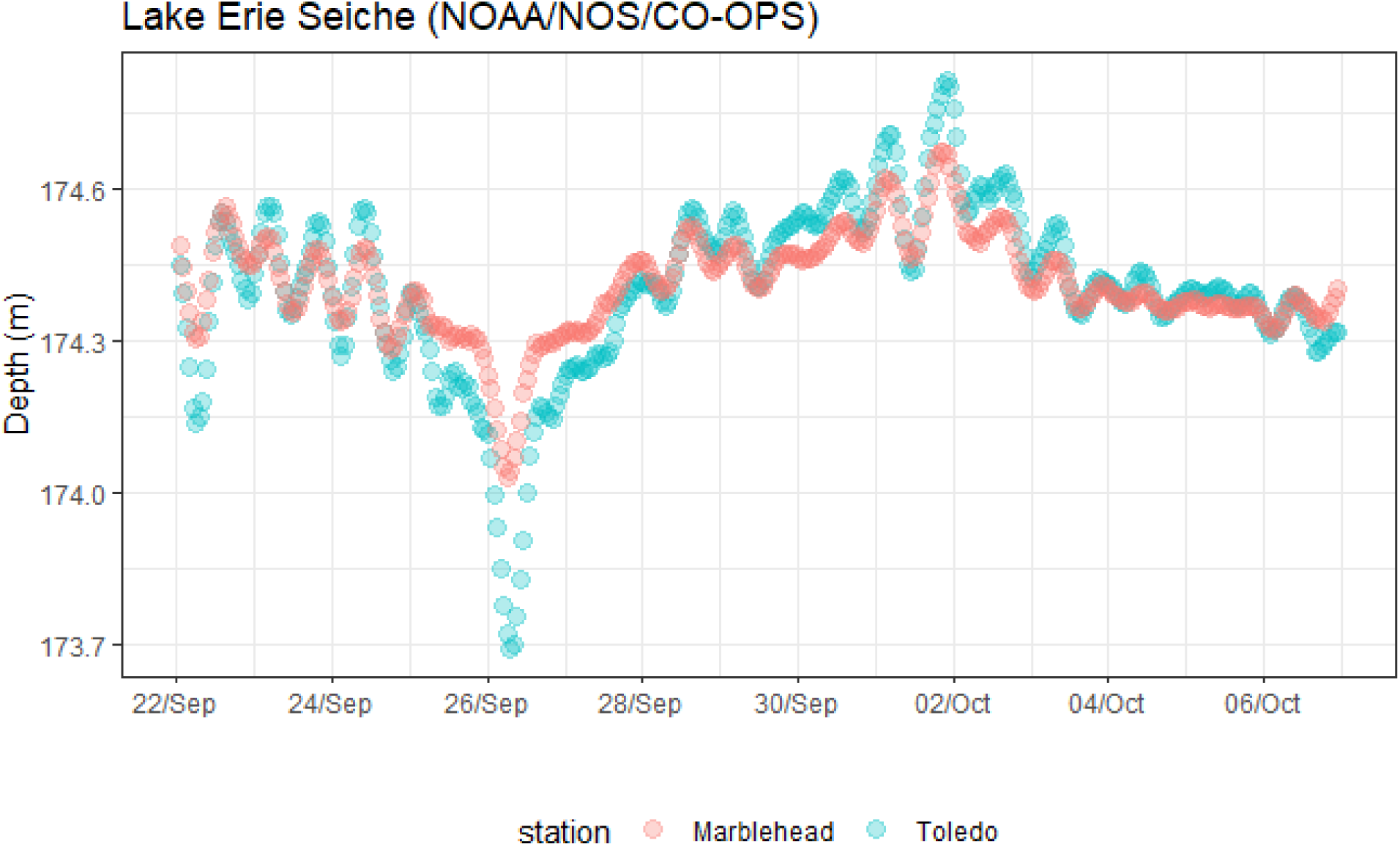
Lake Erie water level during a storm surge (seiche event). Colors are about two stations. Toledo station in blue is west of Crane Creek site and Marblehead is in between sites Portage River and Old Woman Creek.

## REFERENCES

1. Aeppli, M., A. Thompson, C. Dewey & S. Fendorf, 2022. Redox properties of solid phase electron acceptors affect anaerobic microbial respiration under oxygen-limited conditions in floodplain soils. Environmental Science & Technology 56(23):17462–17470.

2. AGIUSA, 2005. Instruction Manual. Austin, TX, USA.

3. Burgin, A. & T. Loecke, 2023. The biogeochemical redox paradox: how can we make a foundational concept more predictive of biogeochemical state changes? Biogeochemistry:1–22.

4. Burri, N. M., R. Weatherl, C. Moeck & M. Schirmer, 2019. A review of threats to groundwater quality in the anthropocene. Science of the Total Environment 684:136–154.

5. Condon, L. E., S. Kollet, M. F. Bierkens, G. E. Fogg, R. M. Maxwell, M. C. Hill, H. J. H. Fransen, A. Verhoef, A. F. Van Loon & M. Sulis, 2021. Global groundwater modeling and monitoring: Opportunities and challenges. Water Resources Research 57(12):e2020WR029500.

6. Fan, D., Y. Lan, P. G. Tratnyek, R. L. Johnson, J. Filip, D. M. O’Carroll, A. Nunez Garcia & A. Agrawal, 2017. Sulfidation of iron-based materials: a review of processes and implications for water treatment and remediation. Environmental science & technology 51(22):13070–13085.

7. Farhadzadeh, A., M. R. Hashemi & S. Neill, 2017. Characterizing the Great Lakes hydrokinetic renewable energy resource: Lake Erie wave, surge and seiche characteristics. Energy 128:661–675.

8. Fiedler, S. & M. Sommer, 2000. Methane emissions, groundwater levels and redox potentials of common wetland soils in a temperate_Jhumid climate. Global Biogeochemical Cycles 14(4):1081–1093.

9. Frohne, T., J. Rinklebe, R. A. Diaz-Bone & G. Du Laing, 2011. Controlled variation of redox conditions in a floodplain soil: Impact on metal mobilization and biomethylation of arsenic and antimony. Geoderma 160(3-4):414–424.

10. Fulda, B., A. Voegelin & R. Kretzschmar, 2013. Redox-controlled changes in cadmium solubility and solid-phase speciation in a paddy soil as affected by reducible sulfate and copper. Environmental Science & Technology 47(22):12775–12783.

11. Gómez-Gener, L., A. R. Siebers, M. I. Arce, S. Arnon, S. Bernal, R. Bolpagni, T. Datry, G. Gionchetta, H.-P. Grossart & C. Mendoza-Lera, 2021. Towards an improved understanding of biogeochemical processes across surface-groundwater interactions in intermittent rivers and ephemeral streams. Earth-Science Reviews 220:103724.

12. Herdendorf, C. E., 1992. Lake Erie coastal wetlands: an overview. Journal of Great Lakes Research 18(4):533–551.

13. Honma, T., H. Ohba, A. Kaneko-Kadokura, T. Makino, K. Nakamura & H. Katou, 2016. Optimal soil Eh, pH, and water management for simultaneously minimizing arsenic and cadmium concentrations in rice grains. Environmental Science & Technology 50(8):4178–4185.

14. Indivero, J., A. N. Myers-Pigg & N. D. Ward, 2021. Seasonal Changes in the Drivers of Water Physico-Chemistry Variability of a Small Freshwater Tidal River. Frontiers in Marine Science 8:607664.

15. Jia, H., Y. Shi, X. Nie, S. Zhao, T. Wang & V. K. Sharma, 2020. Persistent free radicals in humin under redox conditions and their impact in transforming polycyclic aromatic hydrocarbons. Frontiers of Environmental Science & Engineering 14:1–11.

16. Kappler, A., C. Bryce, M. Mansor, U. Lueder, J. M. Byrne & E. D. Swanner, 2021. An evolving view on biogeochemical cycling of iron. Nature Reviews Microbiology 19(6):360–374.

17. Kim, K. H., J. W. Heiss, X. Geng & H. A. Michael, 2020. Modeling hydrologic controls on particulate organic carbon contributions to beach aquifer biogeochemical reactivity. Water Resources Research 56(10):e2020WR027306.

18. Klump, S., O. A. Cirpka, H. Surbeck & R. Kipfer, 2008. Experimental and numerical studies on excess_Jair formation in quasi_Jsaturated porous media. Water resources research 44(5).

19. Kohfahl, C., G. Massmann & A. Pekdeger, 2009. Sources of oxygen flux in groundwater during induced bank filtration at a site in Berlin, Germany. Hydrogeol J 17(3):571–578.

20. Kowalski, K. P., D. A. Wilcox & M. J. Wiley, 2009. Stimulating a Great Lakes coastal wetland seed bank using portable cofferdams: implications for habitat rehabilitation. Journal of Great Lakes Research 35(2):206–214.

21. Kumar, A. R. & P. Riyazuddin, 2012. Seasonal variation of redox species and redox potentials in shallow groundwater: a comparison of measured and calculated redox potentials. Journal of Hydrology 444:187–198.

22. Li, Y., Y. Liu, L. Feng & L. Zhang, 2023. A review: Manganese-driven bioprocess for simultaneous removal of nitrogen and organic contaminants from polluted waters. Chemosphere:137655.

23. Loke, M. H., 1999. Electrical imaging surveys for environmental and engineering studies. A practical guide to 2:70.

24. McMahon, P. & F. Chapelle, 2008. Redox processes and water quality of selected principal aquifer systems. Groundwater 46(2):259–271.

25. Megonigal, J. P., W. Patrick Jr & S. Faulkner, 1993. Wetland identification in seasonally flooded forest soils: soil morphology and redox dynamics. Soil Science Society of America Journal 57(1):140–149.

26. Megonigal, J. P. & M. Rabenhorst, 2013. Reduction–oxidation potential and oxygen. Methods in biogeochemistry of wetlands 10:71–85.

27. Meng, L., R. Zuo, J.-s. Wang, Q. Li, C. Du, X. Liu & M. Chen, 2021. Response of the redox species and indigenous microbial community to seasonal groundwater fluctuation from a typical riverbank filtration site in Northeast China. Ecological Engineering 159:106099.

28. Neogi, S., P. Dash, P. Bhattacharyya, S. Padhy, K. Roy & A. Nayak, 2020. Partitioning of total soil respiration into root, rhizosphere and basal-soil CO2 fluxes in contrasting rice production systems. Soil Research 58(6):592–601.

29. Nimick, D. A., C. H. Gammons & S. R. Parker, 2011. Diel biogeochemical processes and their effect on the aqueous chemistry of streams: A review. Chemical Geology 283(1-2):3–17.

30. NOAA, 2023. National Oceanic and Atmospheric Administration – https://tidesandcurrents.noaa.gov.

31. Noyce, G. L., A. J. Smith, M. L. Kirwan, R. L. Rich & J. P. Megonigal, 2023. Oxygen priming induced by elevated CO2 reduces carbon accumulation and methane emissions in coastal wetlands. Nature Geoscience:1–6.

32. O’Connor, A. E., J. L. Krask, E. A. Canuel & A. J. Beck, 2018. Seasonality of major redox constituents in a shallow subterranean estuary. Geochimica et Cosmochimica Acta 224:344–361.

33. Olid, C., V. Rodellas, G. Rocher-Ros, J. Garcia-Orellana, M. Diego-Feliu, A. Alorda-Kleinglass, D. Bastviken & J. Karlsson, 2022. Groundwater discharge as a driver of methane emissions from Arctic lakes. Nature communications 13(1):3667.

34. Peiffer, S., A. Kappler, S. Haderlein, C. Schmidt, J. Byrne, S. Kleindienst, C. Vogt, H. Richnow, M. Obst & L. T. Angenent, 2021. A biogeochemical–hydrological framework for the role of redox-active compounds in aquatic systems. Nature Geoscience 14(5):264–272.

35. Peixoto, R. B., F. Machado-Silva, H. Marotta, A. Enrich-Prast & D. Bastviken, 2015. Spatial versus day-to-day within-lake variability in tropical floodplain lake CH4 emissions– Developing optimized approaches to representative flux measurements. PLoS One 10(4):e0123319.

36. Pi, K., P. Van Cappellen, L. Tong, Y. Gan & Y. Wang, 2023. Loss of Selenium from Mollisol Paddy Wetlands of Cold Regions: Insights from Flow-through Reactor Experiments and Process-Based Modeling. Environmental Science & Technology 57(15):6228–6237.

37. Regier, P., N. D. Ward, J. Indivero, C. Wiese Moore, M. Norwood & A. Myers_JPigg, 2021. Biogeochemical control points of connectivity between a tidal creek and its floodplain. Limnology and Oceanography Letters 6(3):134–142.

38. Rezanezhad, F., R.-M. Couture, R. Kovac, D. O’connell & P. Van Cappellen, 2014. Water table fluctuations and soil biogeochemistry: An experimental approach using an automated soil column system. Journal of Hydrology 509:245–256.

39. Richards, L. A., R. Kumari, N. Parashar, A. Kumar, C. Lu, G. Wilson, D. Lapworth, V. J. Niasar, A. Ghosh & B. Chakravorty, 2022. Environmental tracers and groundwater residence time indicators reveal controls of arsenic accumulation rates beneath a rapidly developing urban area in Patna, India. Journal of Contaminant Hydrology 249:104043.

40. Rosentreter, J. A., A. V. Borges, B. R. Deemer, M. A. Holgerson, S. Liu, C. Song, J. Melack, P. A. Raymond, C. M. Duarte & G. H. Allen, 2021. Half of global methane emissions come from highly variable aquatic ecosystem sources. Nature Geoscience 14(4):225–230.

41. Santos, I. R., X. Chen, A. L. Lecher, A. H. Sawyer, N. Moosdorf, V. Rodellas, J. Tamborski, H.-M. Cho, N. Dimova & R. Sugimoto, 2021. Submarine groundwater discharge impacts on coastal nutrient biogeochemistry. Nature Reviews Earth & Environment 2(5):307–323.

42. Scheffer, M., S. Carpenter, J. A. Foley, C. Folke & B. Walker, 2001. Catastrophic shifts in ecosystems. Nature 413(6856):591–596.

43. Swatridge, L. L., R. P. Mulligan, L. Boegman, S. Shan & R. Valipour, 2022. Coupled modelling of storm surge, circulation and surface waves in a large stratified lake. Journal of Great Lakes Research 48(6):1520–1535.

44. USGS, 2023. United States Geological Survey. The Quaternary Geologic Map of the Lake Erie. usgs.gov In.

57. Wang, J., D. Halder, L. Wegner, L. Bru□genwirth, J. r. Schaller, M. Martin, D. Said-Pullicino, M. Romani & B. Planer-Friedrich, 2020. Redox dependence of thioarsenate occurrence in paddy soils and the rice rhizosphere. Environmental Science & Technology 54(7):3940–3950.

45. Wang, Y. & N. Chen, 2021. Recent progress in coupled surface–ground water models and their potential in watershed hydro-biogeochemical studies: A review. Watershed Ecology and the Environment 3:17–29.

46. Wanner, P., R. Aravena, J. Fernandes, M. BenIsrael, E. A. Haack, D. T. Tsao, K. E. Dunfield & B. L. Parker, 2019. Assessing toluene biodegradation under temporally varying redox conditions in a fractured bedrock aquifer using stable isotope methods. Water research 165:114986.

47. Ward, N. D., J. P. Megonigal, B. Bond-Lamberty, V. L. Bailey, D. Butman, E. A. Canuel, H. Diefenderfer, N. K. Ganju, M. A. Goñi & E. B. Graham, 2020. Representing the function and sensitivity of coastal interfaces in Earth system models. Nature communications 11(1):2458.

48. Yazbeck, T. & G. Bohrer, 2023. Uncertainties in wetland methane-flux estimates. Global Change Biology.

49. Yu, K., S. P. Faulkner & W. H. Patrick Jr, 2006. Redox potential characterization and soil greenhouse gas concentration across a hydrological gradient in a Gulf coast forest. Chemosphere 62(6):905–914.

50. Yu, X., J. J. LeMonte, J. Li, J. W. Stuckey, D. L. Sparks, J. G. Cargill, C. J. Russoniello & H. A. Michael, 2022. Hydrologic Control on Arsenic Cycling at the Groundwater–Surface Water Interface of a Tidal Channel. Environmental Science & Technology.

51. Zakem, E. J., M. F. Polz & M. J. Follows, 2020. Redox-informed models of global biogeochemical cycles. Nature communications 11(1):5680.

52. Zhang, L., X. Ma, Q. Li, H. Cui, K. Shi, H. Wang, Y. Zhang, S. Gao, Z. Li & A.-J. Wang, 2023. Complementary Biotransformation of Antimicrobial Triclocarban Obviously Mitigates Nitrous Oxide Emission toward Sustainable Microbial Denitrification. Environmental Science & Technology 57(19):7490–7502.

53. Zhang, S. & N. J. Planavsky, 2020. Revisiting groundwater carbon fluxes to the ocean with implications for the carbon cycle. Geology 48(1):67–71.

54. Zhang, Z. & A. Furman, 2021. Soil redox dynamics under dynamic hydrologic regimes-A review. Science of The Total Environment 763:143026.

55. Zhao, G., M. Tan, B. Wu, X. Zheng, R. Xiong, B. Chen, A. Kappler & C. Chu, 2023. Redox Oscillations Activate Thermodynamically Stable Iron Minerals for Enhanced Reactive Oxygen Species Production. Environmental Science & Technology.

56. Zuo, R., M. Pan, J. Li, L. Meng, J. Yang, Y. Zhai, Z. Xue, J. Liu, J. Shi & Y. Teng, 2021. Biogeochemical transformation processes of iron, manganese, ammonium under coexisting conditions in groundwater based on experimental data. Journal of Hydrology 603:127120.

