## Supplementary material for "Groundwater redox dynamics across the terrestrial-aquatic interface of Lake Erie coastal ecosystems": Figure S1

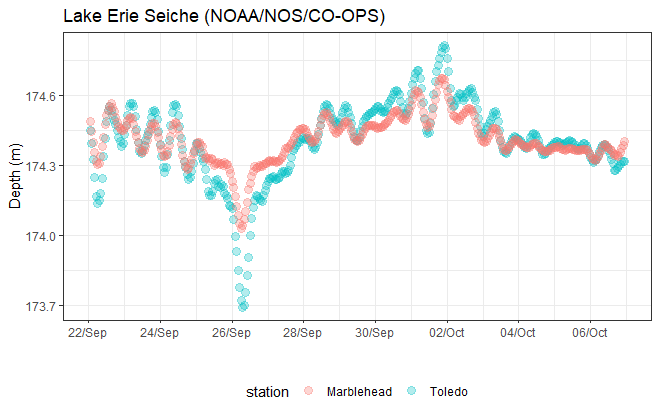

Figure S2. Lake Erie water level during a storm surge (seiche event). Colors are about two stations. Toledo station in blue is west of Crane Creek site and Marblehead is in between sites Portage River and Old Woman Creek.
